# Experimental study of tissue-engineered urethra with collagen / chitosan composite as scaffolds

**DOI:** 10.1101/2020.04.30.070284

**Authors:** Zhai Hongfeng, Qiu Changhong, Jin Jun, Shao Xin

## Abstract

In this article we investigated the preparation of tissue-engineered urethra by using the urethral epithelial subculture cells of male New Zealand young rabbits. We inoculated the epithelial cells of urinary mucosa of male New Zealand young rabbits on collagen, chitosan and collagen chitosan composite as scaffolds to prepare tissue-engineered urethra. The results of inverted phase contrast microscope, HE staining and scanning electron microscope of three kinds of tissue-engineered urethra were compared. What’s more, we reported a new method for quantitative and rapid detection of epithelial cell activity of urinary mucosa in situ by Interactive Laser Cytometer. The collagen chitosan composite was more similar to the extracellular matrix of mammalian. Its three-dimensional porous structure had a high area volume ratio, which was conducive to cell adhesion, growth and metabolism. In vitro, the urethral epithelial cells had been cultured on collagen chitosan composite, and the tissue-engineered urethra was successfully prepared, which laid a solid foundation for further in vivo experiments.

## Introduction

Urethral reconstruction continues to be a challenging field for urologists. Urethral stricture is a disease characterized by a pathological narrowing of the urethra. Treatment for this condition often requires surgery using autologous grafts (urethroplasty). It is common practice to use patient’s own tissue like genital and extragenital skin, bladder and buccal mucosa as a source of the graft [1]. Whilst for some conditions only one or few procedures are standard, over 300 techniques are known for urethral stricture repair [2]. This diversity illustrates the complexity of these conditions and also indicates the lack of one perfect procedure. However, all of these substitutes have limitations compared to the autologous urethral tissue, which can lead to complications (e.g. stricture formation, graft failure). Also, the amount of tissue that can be harvested from a donor site is limited; especially in the case of long defects, this could pose a problem [3]. To overcome these problems, alternative materials for urethral reconstruction have been explored.

In the field of regenerative medicine, tissue engineering (TE) is defined as “*an interdisciplinary field that applies the principles of engineering and life sciences toward the development of biological substitutes that restore, maintain, or improve tissue function or a whole organ*” [4]. As early as the 1980s the first steps were made in culturing urothelial cells. Initially, these cultures were used as an *in vitro* system to study the effects of exogenous substances on tissue [5]. When TE started to evolve, the aim of culturing tissues changed to the replacement of damaged or absent organs. The rationale behind this latter strategy is that, with a limited amount of material (e.g. a small biopsy), a larger graft of autologous cells can be created. Since cells are autologous the problems with rejection are bypassed and, when implanted *in vivo*, the tissue possesses properties similar to those of surrounding tissue.

Alternative and safer approach is to use tissue-engineered graft created in a laboratory using patient’s autologous cells and biocompatible matrix (scaffolds). The choice of biomaterial were mostly acellular matrices derived from natural extracellular matrix. The potential benefit of tissue engineering over the harvesting of autologous tissue is not needing to harvest the tissue, particularly when long lengths of urethra need to be augmented [6].

Although the goal of TE is to give an answer to unmet clinical needs, so far, poor quality results, unclear in animal models, and unrealistic hopes covered the topic [7]. The methods and techniques of isolation, purification and in vitro primary culture and subculture of mammalian urethral epithelial cells have been established in previous experiments [8], and the collagen chitosan composite scaffolds has been prepared with good biocompatibility, biodegradability, plasticity and certain mechanical properties [9]. We have cultured the urethral epithelial subculture cells on collagen chitosan composite to prepare tissue-engineered urethra.

## Materials and Methods

### Tissue engineered urethra with collagen as scaffolds

We took one 8-hole sterile culture plate. The collagen substrate was cut into the bottom of the culture plate (0.28cm^2^) according to the corresponding size, which had been irradiated and sterilized by high-efficiency ultraviolet sterilizer. The urethral epithelial subculture cells of male New Zealand young rabbits were seeded on collagen matrix with a density of 0.7 × 10^4^ / hole. 0.2ml mixed culture medium was added into the culture hole, and cells were cultured in 34 °C, 5% CO_2_ and saturated humidity cell incubator. The growth of the epithelial cells was observed under the inverted phase contrast microscope every day. The culture medium was changed once every 2 days. Cells had been cultured for 7 days.

### Tissue engineered urethra with chitosan as scaffolds

We took one 8-hole sterile culture plate. The chitosan matrix was cut into the bottom of the culture plate (0.28cm2) according to the corresponding size, which had been irradiated and sterilized by high-efficiency ultraviolet sterilizer. The urethral epithelial subculture cells of male New Zealand young rabbits were seeded on chitosan matrix with a density of 0.7×104 / hole. 0.2ml mixed culture medium was added into the culture hole, and cells were cultured in 34 °C, 5% CO2 and saturated humidity cell incubator. The growth of the epithelial cells was observed under the inverted phase contrast microscope every day. The culture medium was changed once every 2 days. Cells had been cultured for 7 days.

### Tissue engineered urethra with collagen chitosan composite as scaffolds

We took one 8-hole sterile culture plate. The collagen chitosan composite matrix was cut into the bottom of the culture plate (0.28cm2) according to the corresponding size, which had been irradiated and sterilized by high-efficiency ultraviolet sterilizer. The urethral epithelial subculture cells of male New Zealand young rabbits were seeded on collagen chitosan composite matrix with a density of 0.7×104 / hole. 0.2ml mixed culture medium was added into the culture hole, and cells were cultured in 34°C, 5% CO2 and saturated humidity cell incubator. The growth of the epithelial cells was observed under the inverted phase contrast microscope every day. The culture medium was changed once every 2 days. The cells had been cultured for 7 days.

### HE staining

Collagen, chitosan and collagen chitosan composite matrix, on which the urethral epithelial cells had been cultured for 7 days, were rinsed twice in PBS solution of pH 7.4, then wrapped with gauze. The samples were put into Carnoy solution for 2 hours, then dehydrated in 85% alcohol. Pathological sections and HE staining were performed.

### CFDA-AM fluorescence staining

CFDA-AM (6-carboxyfluorescein diacetate, acetoxymethyl ester) was dissolved in DMSO to make a storage solution, the concentration of which was 10mg / ml. The solution was refrigerated in dark.

The urethral epithelial cells which had been cultured on collagen chitosan composite scaffolds for 7 days were washed twice with PBS solution (containing Ca^2 +^, Mg^2 +^). CFDA-AM solution was added and the final solution concentration was 10-20 μ g / ml. The cells had been incubated at 37 °C for 10-15 min. After being washed twice with PBS solution (including Ca^2 +^, Mg^2 +^) the cells were observed and taken photograph under fluorescence microscope.

### Ultrastructural observation

Collagen, chitosan and collagen chitosan composite, on which the urethral epithelial cells had been cultured for 7 days, were rinsed twice in PBS solution of pH 7.4, then wrapped in gauze, put into 3% glutaraldehyde phosphate buffer solution, and stored in refrigerator at 4°C for 2 days. Three specimens were dehydrated in 30%、 40%、 50%、 60%、 70%、 80%、 90%、 95% and 100% alcohol for 10 minutes each time, then dried, sprayed gold, and observed with Hitachi 650 SEM.

### Detection by Interactive Laser Cytometer

The urethral epithelial subculture cells (0.7 × 104) of male New Zealand young rabbits had been inoculated on the collagen chitosan composite scaffolds (0.28cm2) for 3 days, 7 days, 14 days and 21 days. Four specimens were stained by double fluorescent CFDA-AM (6-carboxyfluorescein diacetate, acetoxymethyl ester) and PI (Propidium iodide). The culture medium was removed from the culture hole of the cells and collagen chitosan complex. Then 50 μ l CFDA-AM with concentration of 10 μ mol / L, 50 μ l culture medium and 10 μ l PI with concentration of 500 μ g / ml were added and cultured in 37°C incubator for 30 minutes. Then they were washed three times with PBS solution and put on the cover glass for testing.

Quantitative test on computer: the focus plane of each specimen was consistent, magnification was 10 × 2.5, frame was 10, and PMT was 250. Each specimen was measured 10 fields of vision. Each field of vision applied filter lenses with wavelength of 530 nm and 605 nm to measure two values respectively, which represented the fluorescence intensity of CFDA-AM and PI respectively. While the average fluorescence intensity of CFDA represented the number of living cells, the average fluorescence intensity of PI represented the number of dead cells. The data measured in the experiment was analyzed by SAS 9.4 software according to the method of variance analysis.

## Results

### Observation under inverted phase contrast microscope

Collagen, chitosan and collagen chitosan composite as scaffolds had good affinity to urethral epithelial cells, providing a good cell interface, which was conducive to the adhesion and proliferation of urethral epithelial cells. One day after inoculation, the number of epithelial cells increased significantly. The cells grew well on the three substrates and the number of cells increased day after day. Besides, round or quasi round epithelial cells were found on different layers of the focus, which proved that the three-dimensional growth of urethral epithelial cells on the scaffolds. Microscopically, the number of cells on the collagen chitosan composite scaffolds was more than that on the collagen and chitosan scaffolds when the cells were cultured for 7 days. It shows that the collagen chitosan composite scaffolds was more similar to the extracellular matrix of mammalian, and its three-dimensional porous structure had a high area volume ratio, which was conducive to cell adhesion, growth and metabolism.

**Fig 1.**
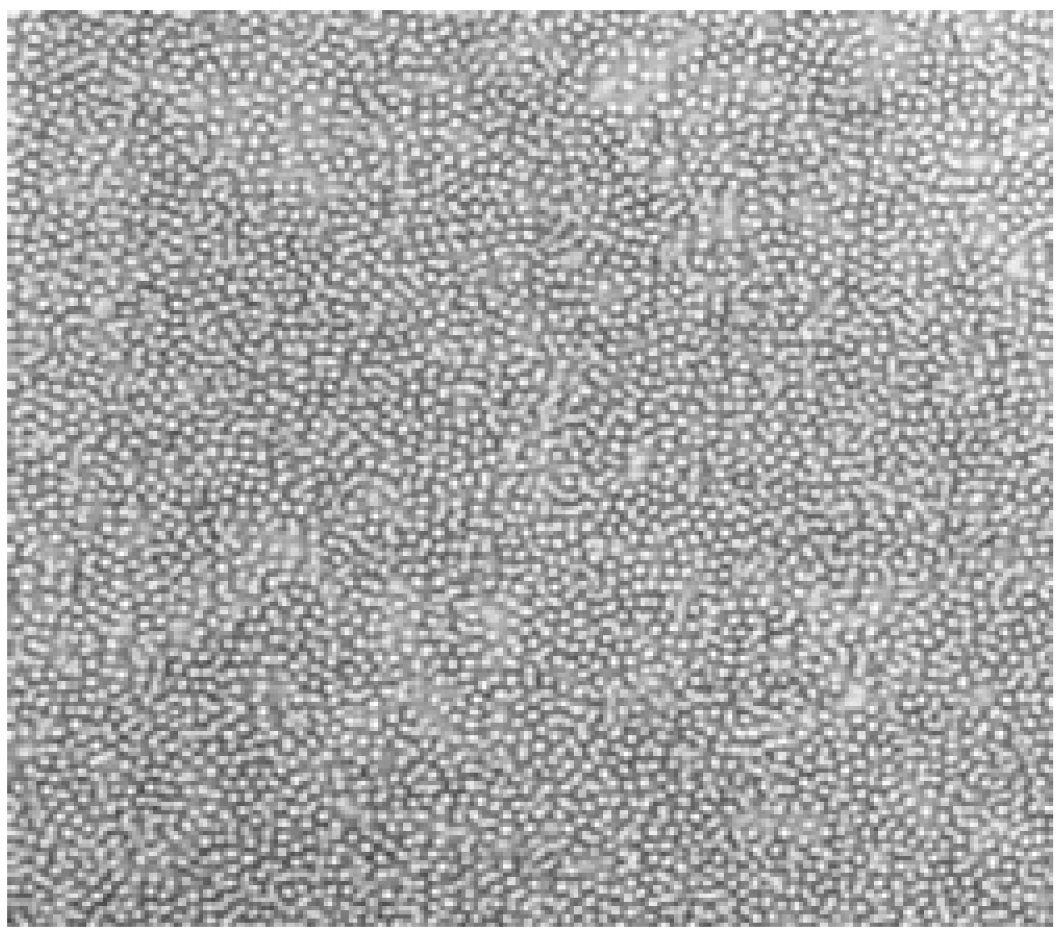
Urethral epithelial cells were cultured on collagen substrate for 7 days (× 100)

**Fig 2.**
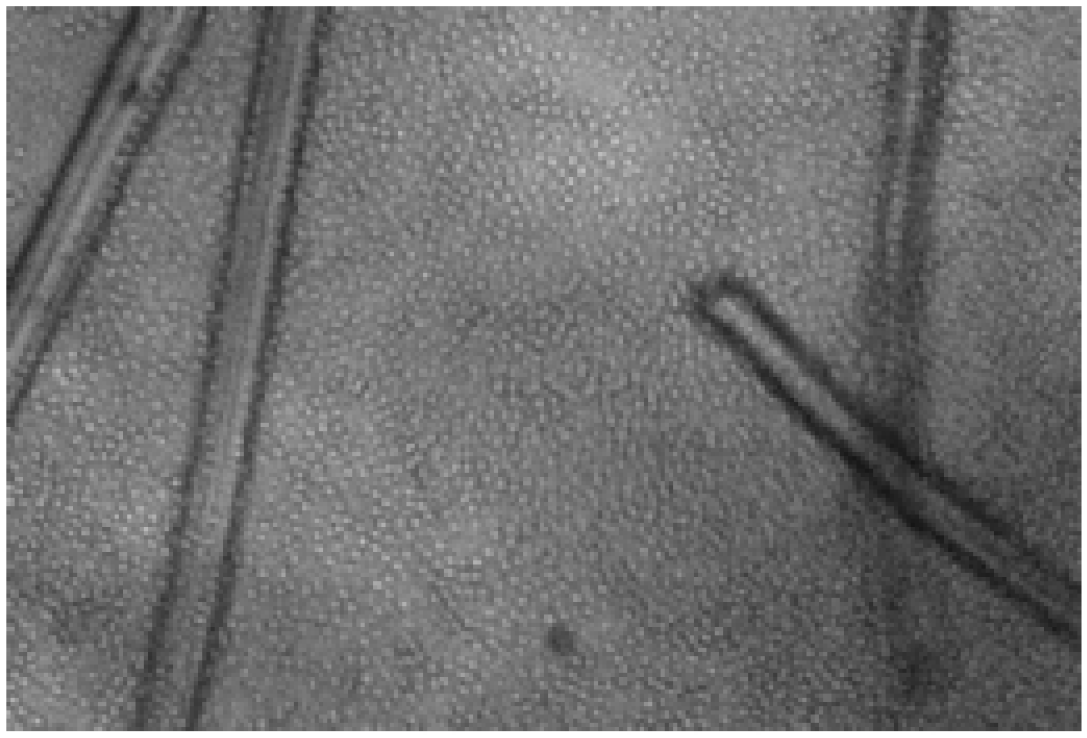
Urethral epithelial cells were cultured on chitosan as scaffolds for 7 days (× 100)

**Fig 3.**
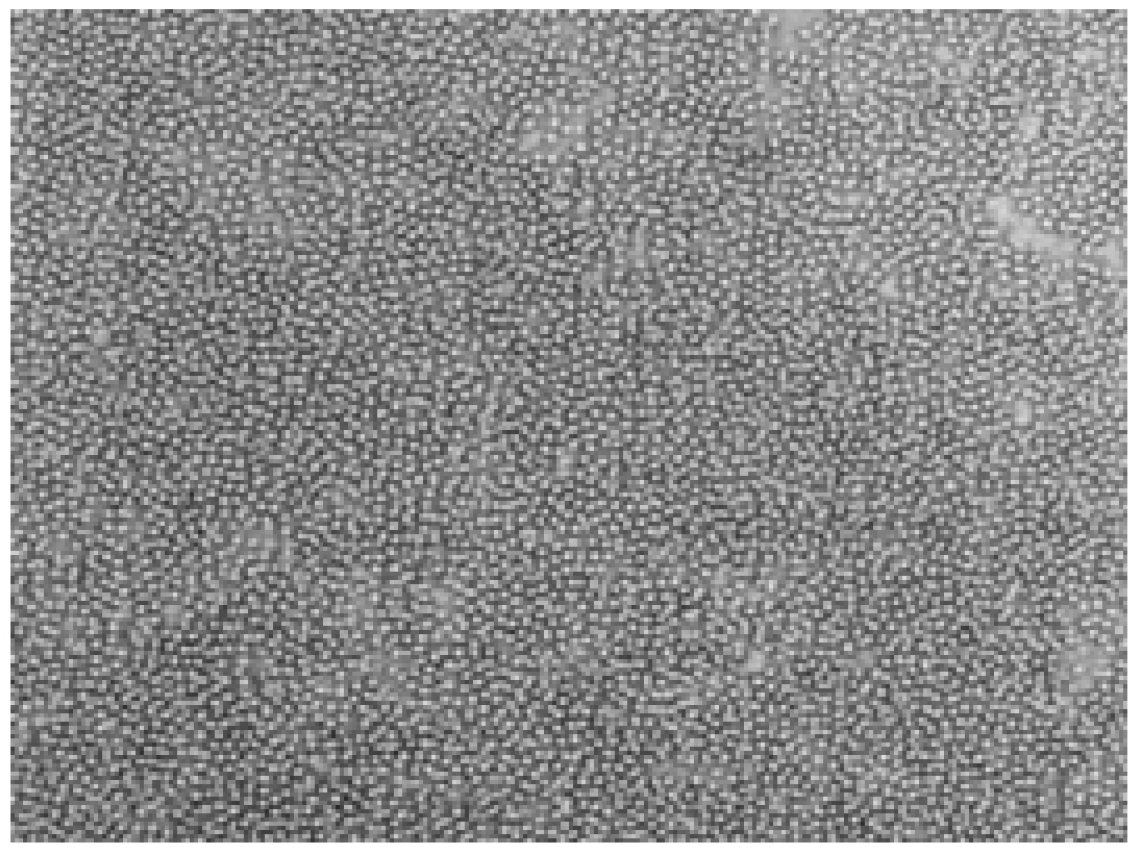
Urethral epithelial cells were cultured on collagen chitosan composite as scaffolds for 7 days (× 100)

**Fig 4.**
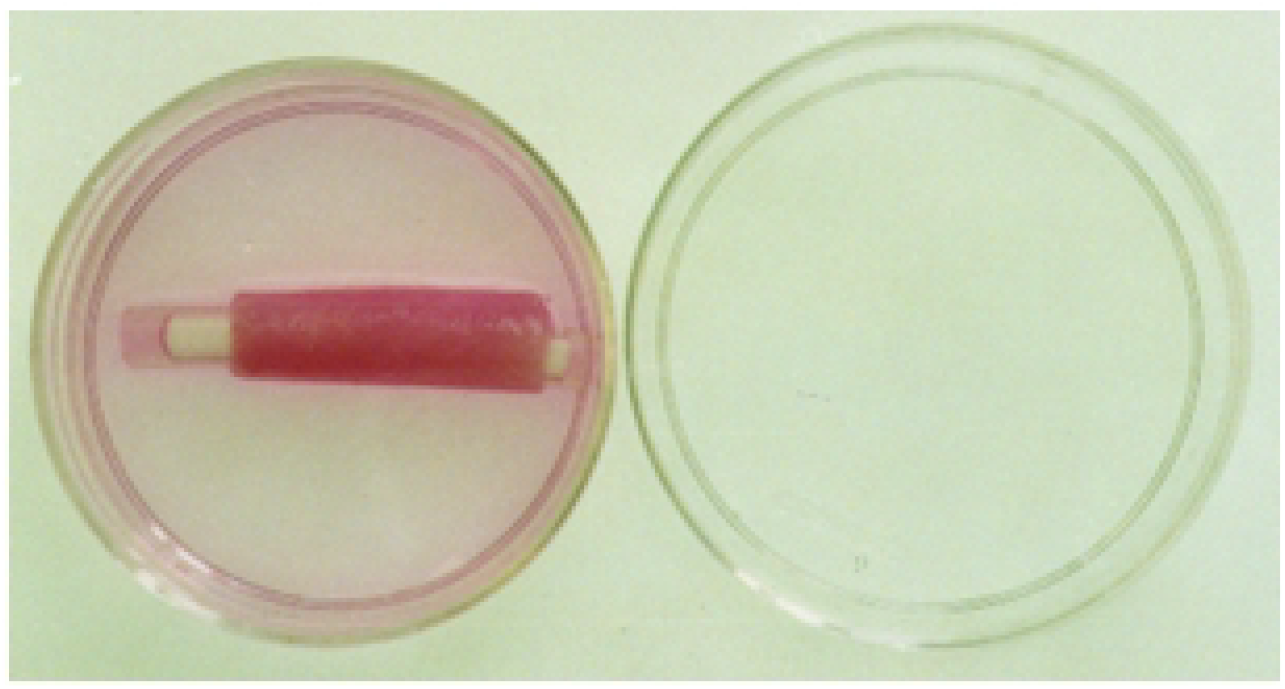
Tissue engineered urethra with collagen chitosan composite as scaffolds

### HE staining

Compared with the monolayer cultured cells, the size of the cells grown on collagen, chitosan and collagen chitosan composite materials was larger, the cells were nearly spherical, full and three-dimensional. The nucleus was dark blue and the cytoplasm was light pink. The density of the urothelial cells grown on the collagen chitosan composite was high.

**Fig 5.**
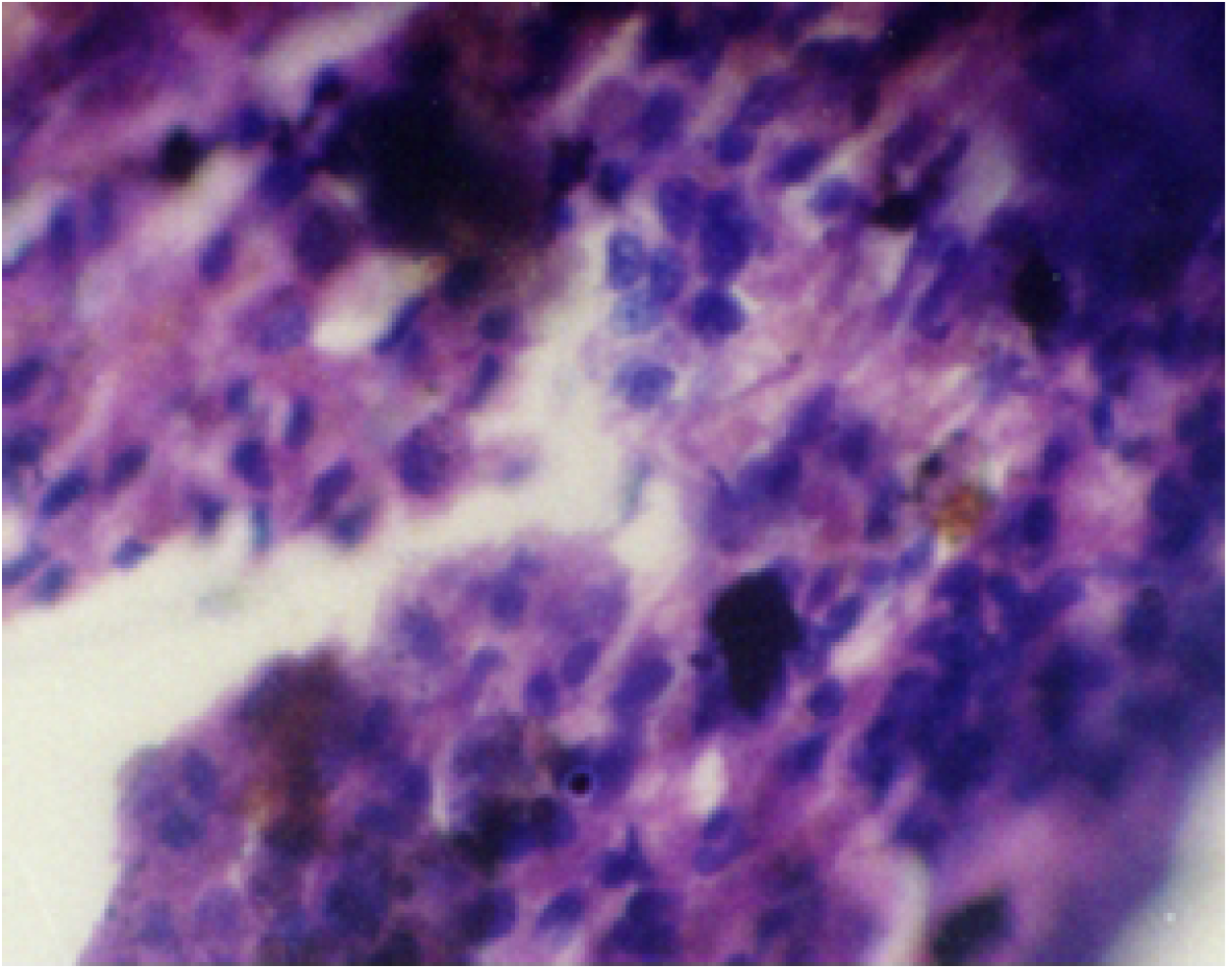
HE staining of tissue-engineered urethra with collagen as scaffolds (× 400)

**Fig 6.**
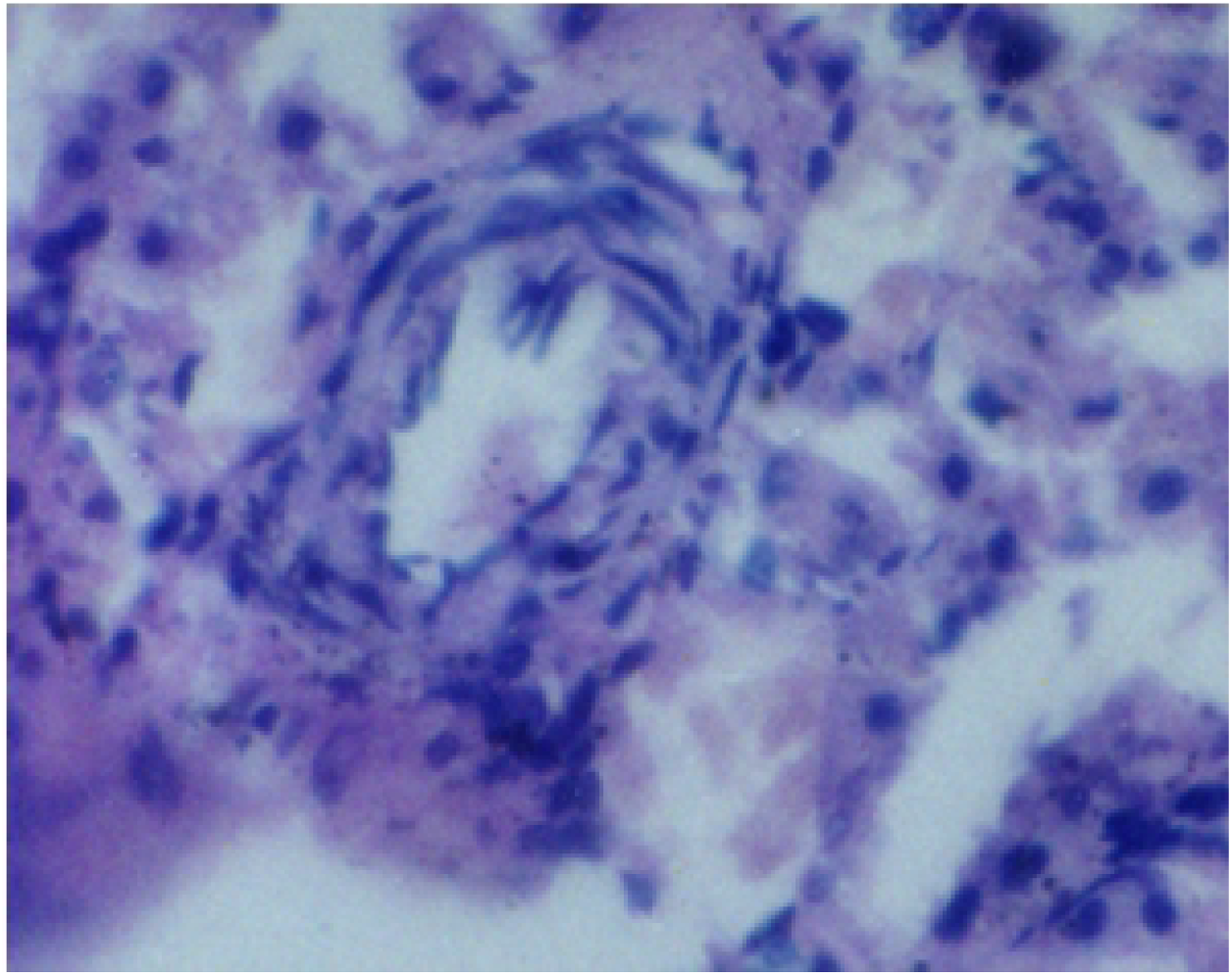
HE staining of tissue-engineered urethra with chitosan as scaffolds (× 400)

**Fig 7.**
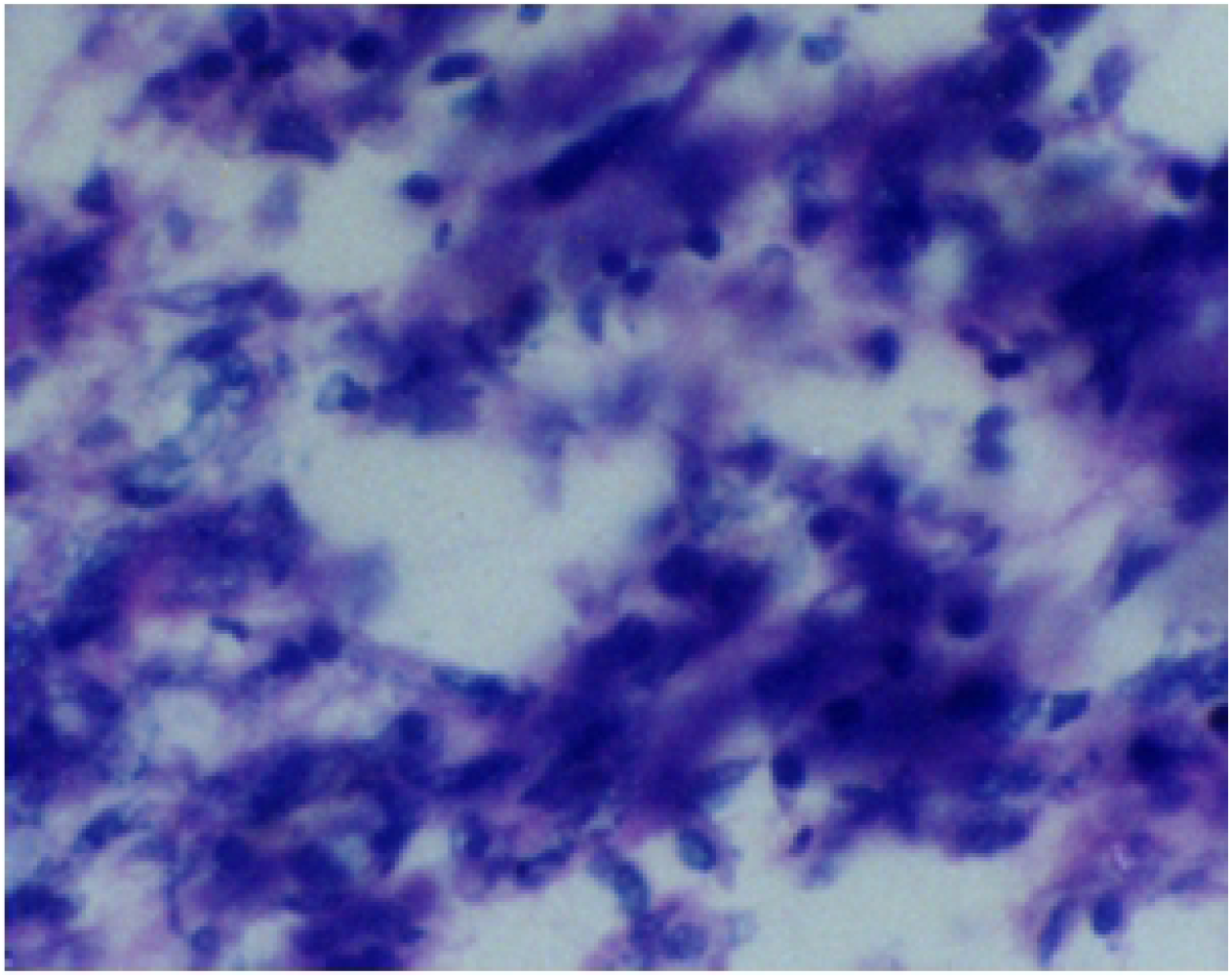
HE staining of tissue-engineered urethra with collagen chitosan composite as scaffolds (× 400)

### Fluorescent staining

The specimen prepared by fluorescent staining must be observed under a special fluorescent microscope. Fluorescent staining is to use specific fluorescent dyes to directly stain the chemical components in the tissue and cells. The advantages of fluorescent staining are high sensitivity, low concentration of dye solution and little damage to cells, which can be used for living staining. CFDA-AM can be absorbed by cells, but it can only be hydrolyzed in living cells to produce CFDA, which makes the cytoplasm produce green fluorescence. On the contrary the dead cells do not.

We observed these components directly under the fluorescent microscope. Under the fluorescence microscope, there were many small round green fluorescent cells on the collagen chitosan composite carrier, which were living urethral epithelial cells stained by CFDA. The results also showed that there were green fluorescent cells in different layers, which indicated that the epithelial cells had three-dimensional growth on the scaffolds.

**Fig 8.**
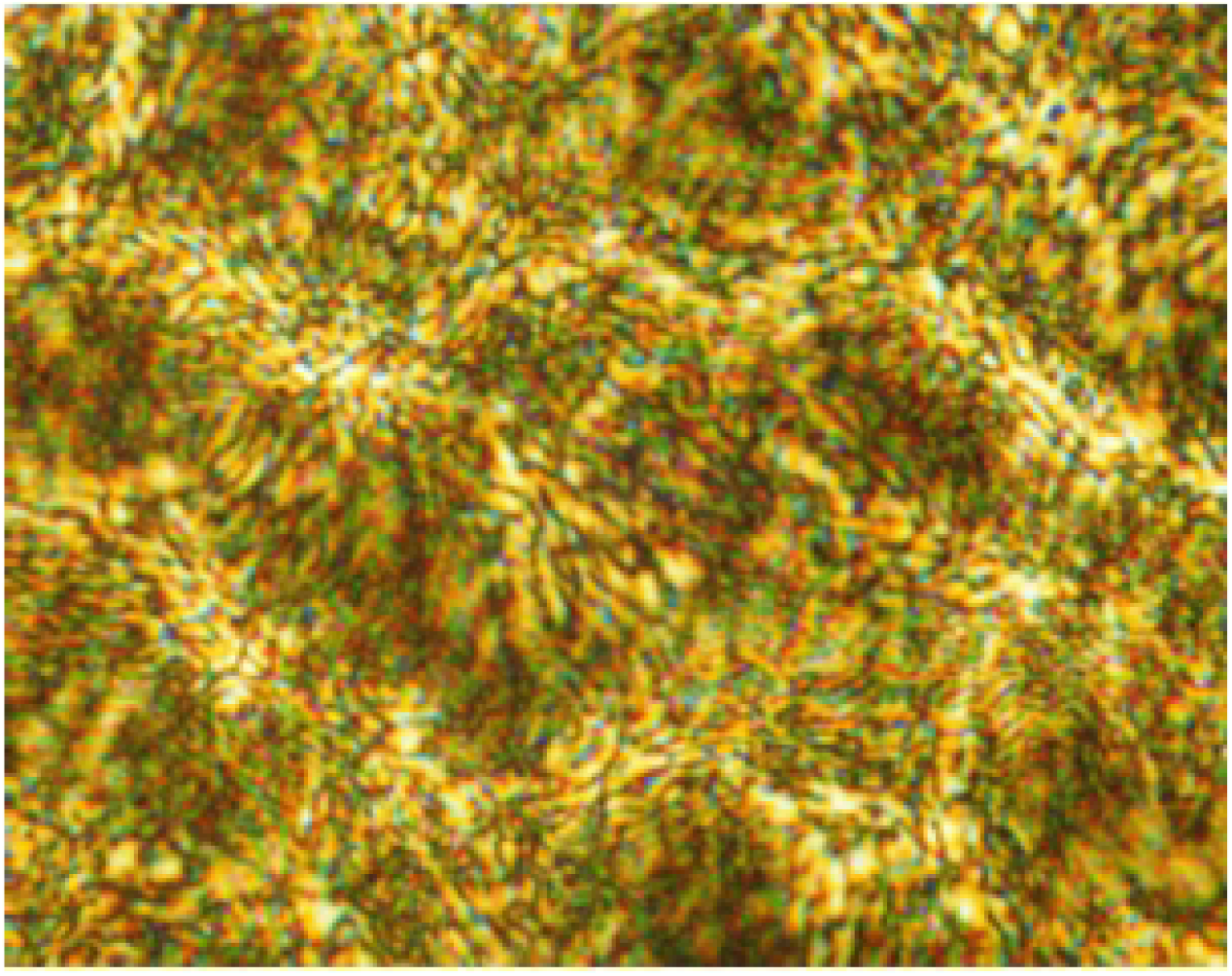
Fluorescence staining (× 100) of urethral epithelial cells which had been cultured for 7 days on collagen chitosan composite as scaffolds

### Ultrastructure

Under the scanning electron microscope, the porosity of collagen and collagen chitosan composite as scaffolds was high, showing three-dimensional network structure. There were many epithelial cells inside and outside the pores. The cells were evenly distributed inside the materials, round or oval, mostly in mosaic arrangement, with microvilli and ridge shape cytoplasmic folds on the surface. However, the number of cells on the chitosan as scaffolds was less, which were also round or oval.

**Fig 9.**
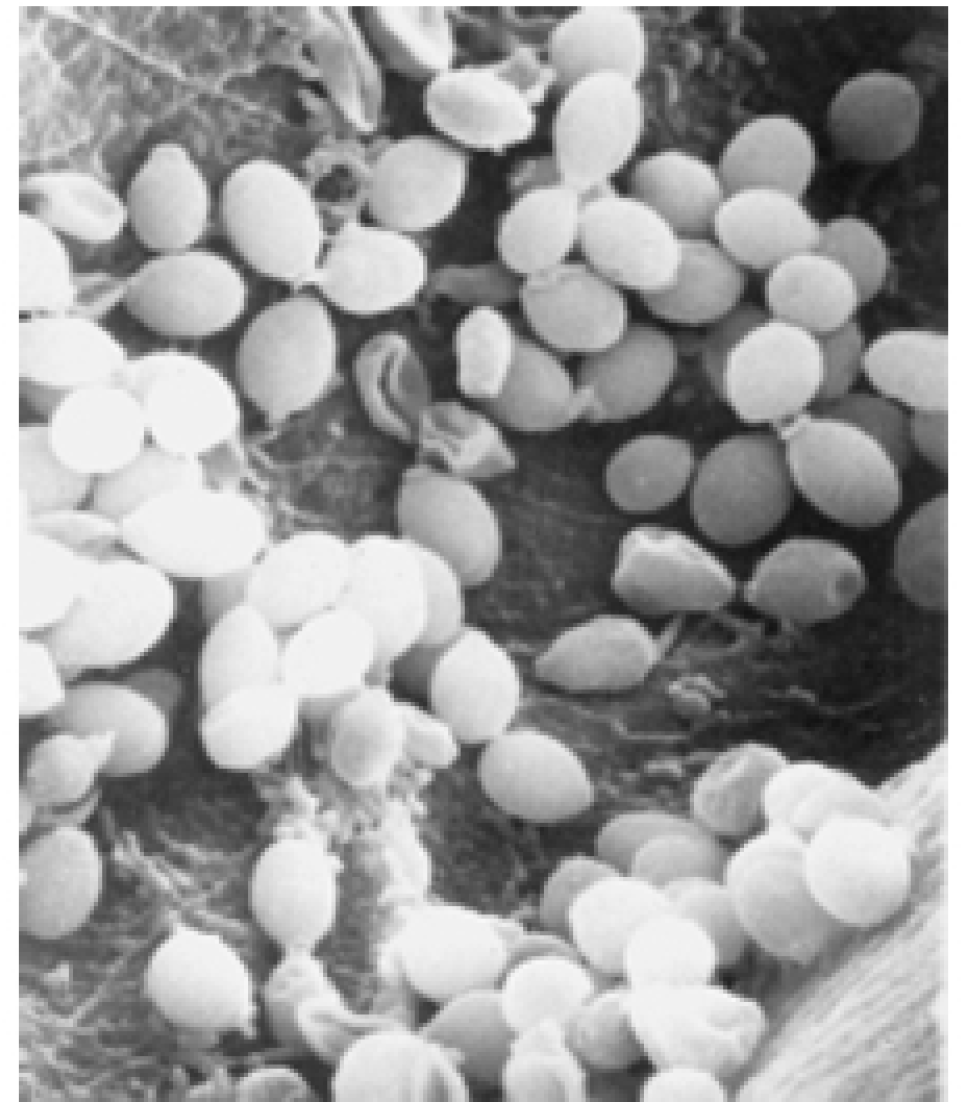
SEM appearance of tissue-engineered urethra with collagen as scaffolds (× 3500)

**Fig 10.**
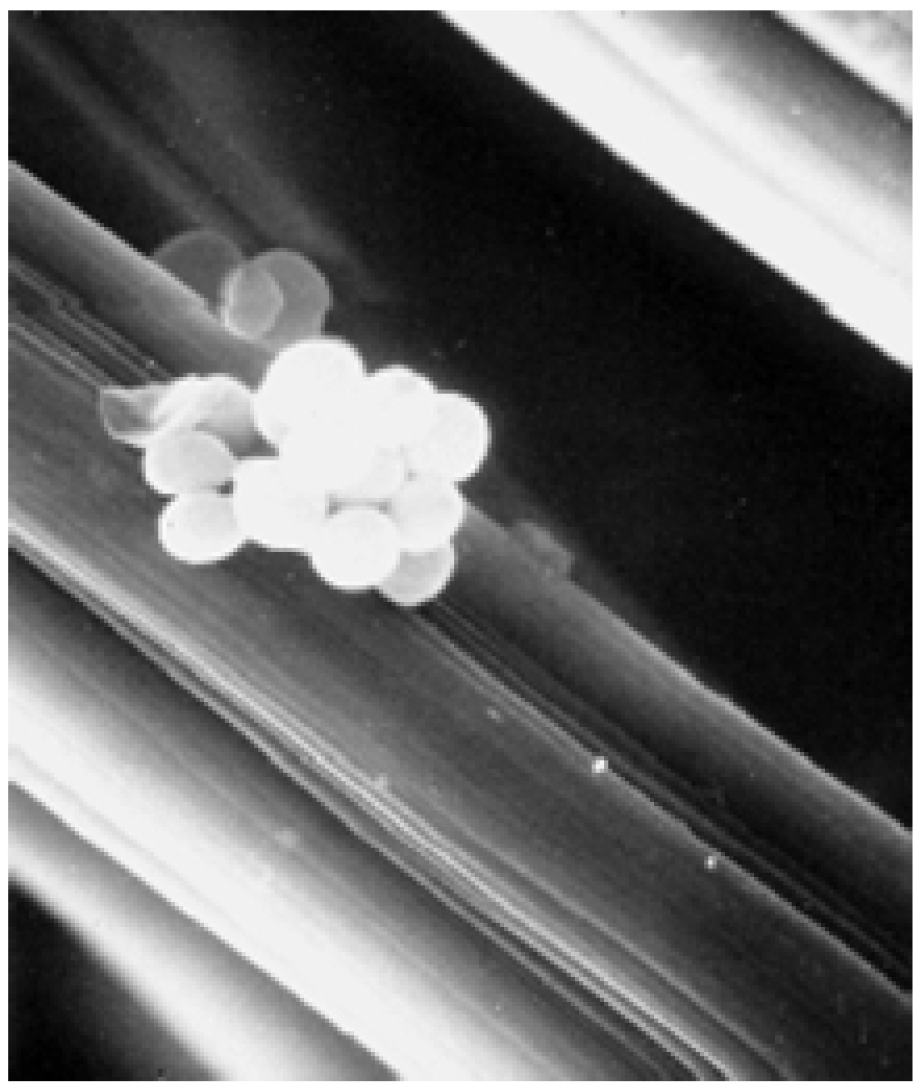
SEM appearance of tissue-engineered urethra with chitosan as scaffolds (× 3500)

**Fig 11.**
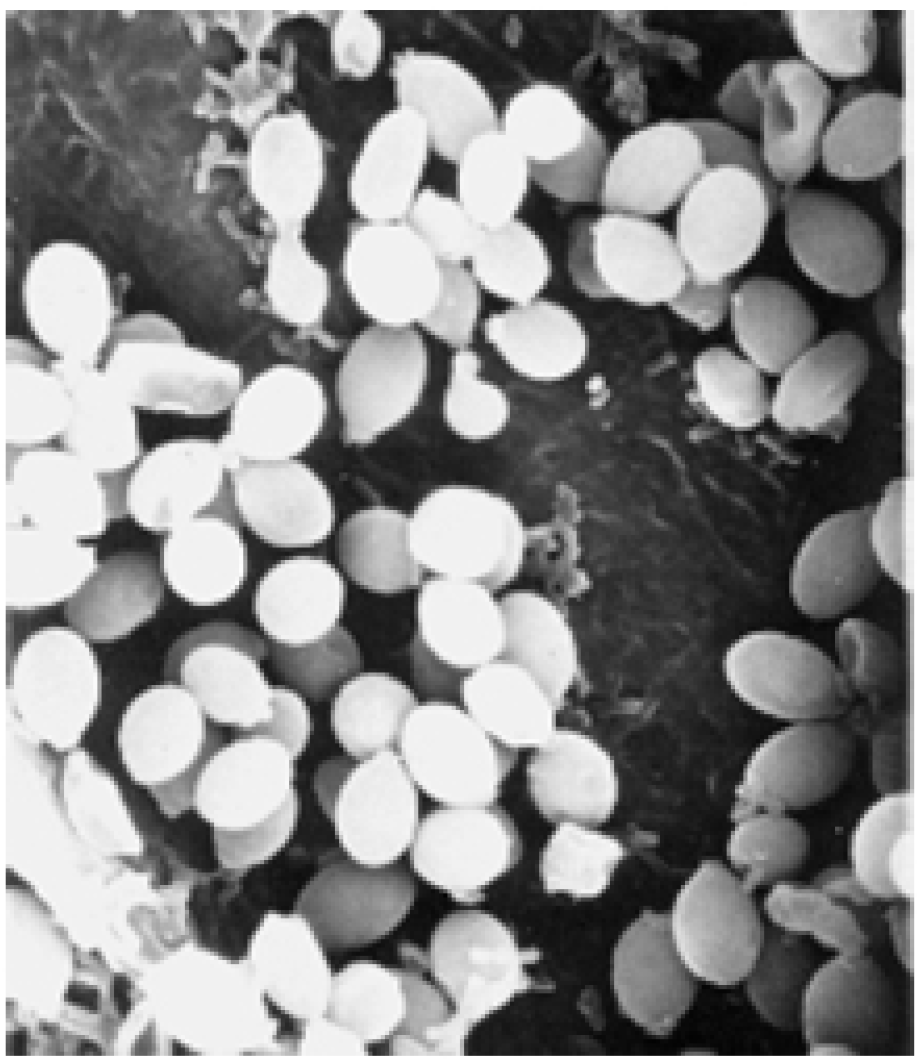
SEM appearance of tissue-engineered urethra with collagen chitosan composite as scaffolds (× 3500)

**Fig 12.**
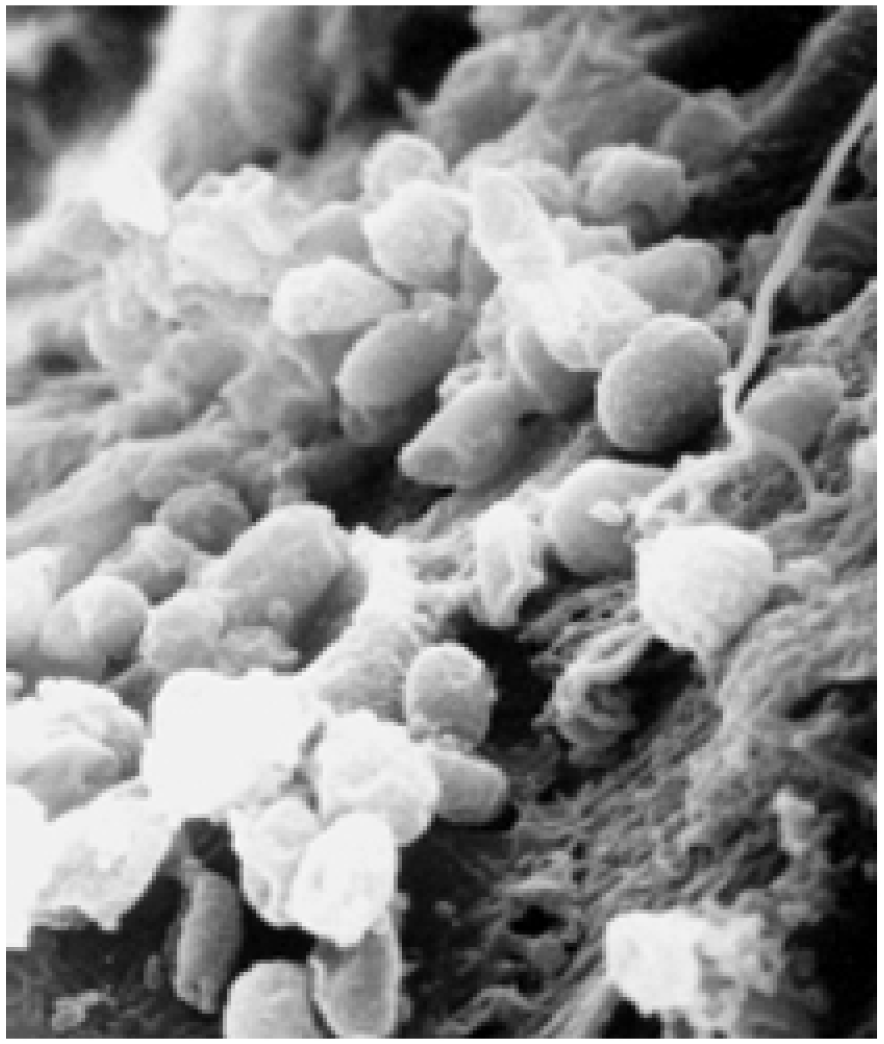
The morphology of cross section scanning of tissue-engineered urethra with collagen chitosan composite as scaffolds showed that the epithelial cells had three-dimensional growth on the scaffolds (× 3500)

**Fig 13.**
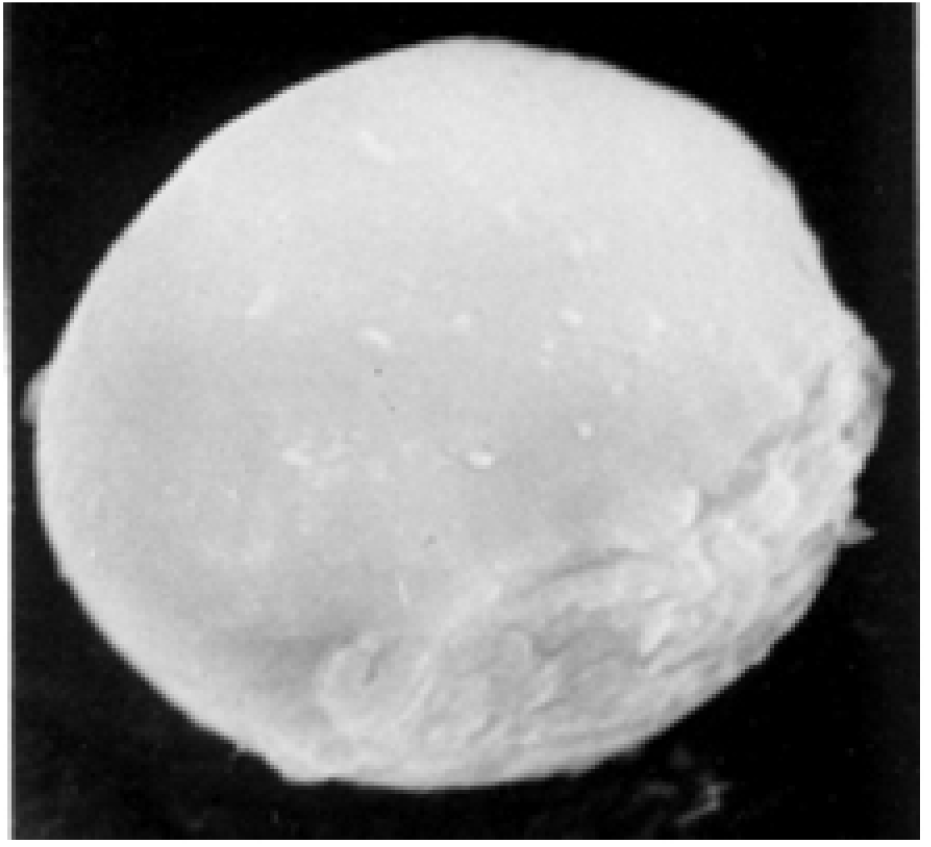
Enlarged scanning electron microscopic appearance of epithelial cells on the collagen chitosan composite as scaffolds (× 10000)

### Detection by Interactive Laser Cytometer

The data measured by Interactive Laser Cytometer were analyzed by SAS 9.4 software according to the method of variance analysis.

**Table 1.** Mean fluorescence intensity of epithelial cells on collagen chitosan composite as scaffolds (mean ± SD)

| Culture time | Mean fluorescence intensity of living cells | Mean fluorescence intensity of dead cells | Fluorescence intensity of living / dead cells |
| --- | --- | --- | --- |
| 3 days | 109360428 ± 13306479 | 32852818 ± 2042098. 0 | 3. 3416 ± 0. 4696 |
| 7 days | 204428078 ± 13094247 | 33401201 ± 1701639. 8 | 6. 1447 ± 0. 6250 |
| 14 days | 55321290 ± 8649102. 4 | 32998420 ± 2660179. 0 | 1. 6791 ± 0. 2319 |
| 21 days | 47018059 ± 3330548. 0 | 34348555 ± 2480254. 7 | 1. 3704 ± 0. 1263 |

**Fig 14.**
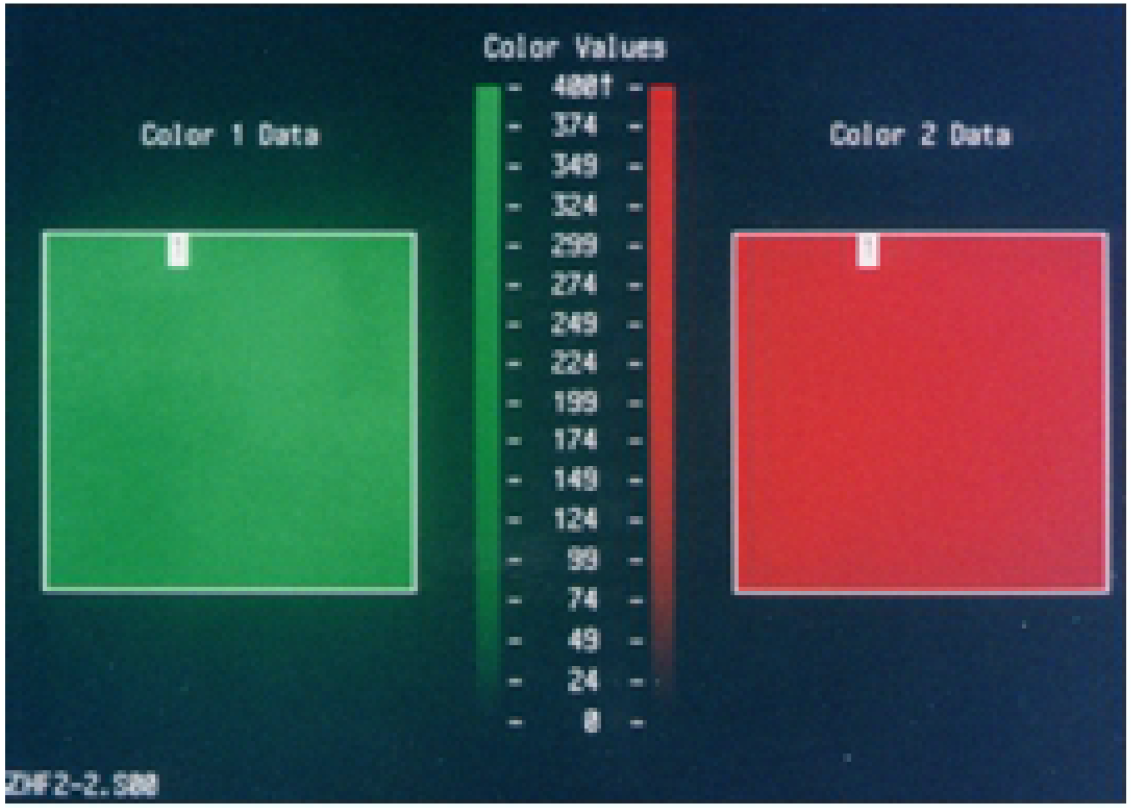
Fluorescence intensity detection by Interactive Laser Cytometer of living cells and dead cells on collagen chitosan composite as scaffolds

From the experimental results above, it showed that the urethral epithelial cells grew well on the collagen chitosan composite as scaffolds, and the number of living cells increased with the culture time. What’s more, the fluorescence intensity of living cells was the highest and the fluorescence intensity of living cells / dead cells was also the highest when the urethral epithelial cells had been cultured for 7 days, indicating that the number of living cells was the highest. After 7 days, both the fluorescence intensity of living cells and the fluorescence intensity of living cells / dead cells decreased rapidly at 14 and 21 days, even lower than that at 3 days. Maybe the cells were affected by density inhibition and contact inhibition. In addition, with the culture time, the fluorescence intensity of dead cells changed little, only slightly increased. This result provided an objective basis for an opportune moment to implant tissue-engineered urethra into animal’s body when the urethral epithelial cells were inoculated on collagen chitosan composite material as scaffolds.

**Fig 15.**
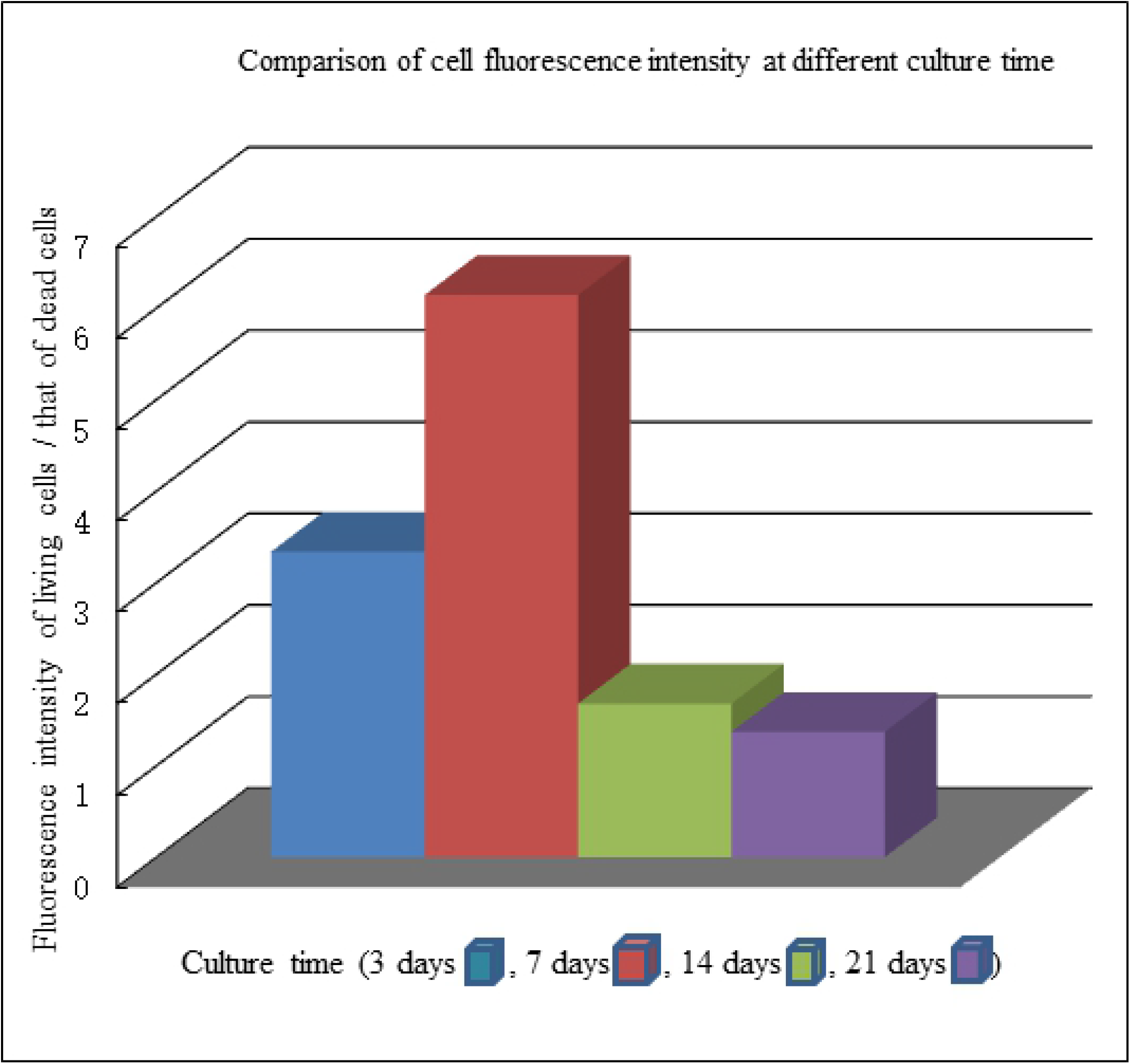
Comparison of fluorescence intensity of living cells and dead cells at different culture time

## Discussion

### Urethral reconstruction using urethral mucosa cells

Several cell types or tissues are of interest for reconstruction: (i) Urethral epithelial lining (urothelium of [pseudo]stratified columnar epithelium) and SMC, because they form the most important layers of the urethra. (ii) Buccal mucosa, as this is often used as a graft in urethroplasties. (iii) Bladder urothelium, as this is easy to expand from small bladder biopsies or can be grown from cells isolated either from bladder washings or urine. (iv) Stem cells from non-urologic tissue (such as adipose tissue) are also under investigation because they are easier to obtain and have the capacity to differentiate to urothelial and myogenic lineages [3].

Although experience with differentiation of stem cells (either isolated from urine or from adipose tissue) towards different lineages is gaining ground, protocols with in vitro expansion of original tissue are better established at this moment. It is noteworthy that no research has been performed with pseudostratified urethral epithelium [3]. The cultured urethral epithelial cells are capable of extensive expansion, and can be passaged 11 to 13 generations in vitro [8].

In the experiment the urethra mucosa epithelial cells of male New Zealand young rabbits were seeded on collagen, chitosan and collagen chitosan composite as scaffolds to prepare tissue-engineered urethra. Morphological examination of inverted phase contrast microscope, HE staining and scanning electron microscopy of the tissue-engineered urethra showed satisfactory results. Perhaps in the future with a small biopsy inside urethral orifice, a larger graft of autologous cells can be created to reconstruct urethral defect.

### Selection of biomimetic cell carrier

When cells are cultured on a scaffold, a choice must be made between a synthetic and a biological scaffold. An advantage of synthetic scaffolds is that the structure and properties can be altered to specific conditions, bearing in mind that the mechanical properties of the scaffold itself influence cellular adherence and proliferation. A disadvantage of classic synthetic scaffolds is the absence of ECM proteins, which are present on biological scaffolds. These proteins are biologically active and play an important role in supporting cellular proliferation and differentiation. The decellularization process of biological scaffolds is important for preserving these ECM proteins [10].

Considerations regarding the choice of scaffold are that biological scaffolds have the advantage of bioactive ECM proteins on their surface and do not require a technologically challenging production process [11]. On the other hand, bioactive synthetic scaffolds loaded with ECM components and growth factors to attract cells and provide a niche as in native tissue, are currently being tested in vitro and in animal models [12].

The composite cell carrier should have good biocompatibility, appropriate mechanical strength, air permeability and moisture permeability, appropriate pore size and channel, which is conducive to cell growth and new tissue and capillary growth, especially with the growth of cells and new tissue. Meanwhile the carrier material itself continues to degrade to achieve synchronization, and the transplanted tissue and autologous tissue melt into each other integration [13]. This is an ideal carrier model. From the point of view of bionics and through systematic biological evaluation, we found that the collagen chitosan composite has good biocompatibility, non-toxic to the body, none mutagenic, strong affinity to cells, and is conducive to cell growth and differentiation. The material has three-dimensional structure, high porosity, suitable for three-dimensional cell growth, and has a certain mechanical strength, which can provide a good carrier for new tissue [3].

Chitosan in the composite carrier is particularly important for its performance. Neither collagen nor chitosan can meet the needs of tissue engineering cell culture considering the strength and degradation rate. But the mechanical strength of the carrier is increased when collagen and chitosan are combined. Chitosan is a straight chain amino polysaccharide, soluble in weak acid, easy to form film, whose mechanical strength is high. When mixed with collagen dispersion to form a membrane, a very thin membrane is formed between the collagen fibers, which enhances the tension between the fibers. When it is wet, it can still keep upright and is easy to operate.

Chitosan delays the degradation of the composite carrier. Collagen fibers degrade rapidly in vivo. Adding chitosan can delay the time of collagen degradation, which is beneficial to the growth of cells and autologous tissue. This is also the characteristic of ideal tissue engineering scaffolds. The degradation of collagen is mainly performed by collagenase, while chitosan is degraded by lysozyme. In vivo, the content of these two enzymes is different. When lysozyme decomposes the chitosan wrapped in the collagen fiber, collagenase can degrade the collagen fiber, which may be the reason why the degradation time is delayed [14].

### Selection of fluorescent probe for determining the viability of living cells

The fluorescent probe has advantages in detecting the viability of cultured cells, because it is more economical than the isotope detection method and has no radiation damage. The colorimetric method has high sensitivity, but that method must be completed within 3-5 minutes. When the commonly used trypan blue colorimetric method is used to determine the viability of cells, the number of dyed blue cells will keep increasing. Therefore, we chose the fluorescent probe as a means to detect the living cells [15]. The CFDA-AM and PI probes we selected have different performance and excitation wavelength. CFDA-AM itself does not produce fluorescence. When it is decomposed by esterase to produce CFDA it produces green fluorescence. Therefore, it is a permeable cell dye. When incubated with cells, CFDA-AM can enter cells through cell membrane and be decomposed by non-specific esterase to produce CFDA. What’s more, many negative charges contained in CFDA-AM are retained in cells and will not leak out of cells [16]. CFDA has been used to determine the activity of a variety of cells, and the results are consistent with ^51^Cr release method [17]. After CFDA-AM staining, the more the number of living cells, the stronger the green fluorescence intensity. Therefore, CFDA-AM is a reliable specific fluorescent probe for detecting living cells.

PI can produce red fluorescence, and it can’t pass through the cell membrane of living cells. PI is not a permeable cell dye. The cell membrane is destroyed after cell death, while PI can pass through the cell membrane and combine with nucleic acid producing red fluorescence. Therefore, PI can only be combined with dead cells. The more the number of dead cells, the stronger the red fluorescence intensity, so it is a reliable specific fluorescent probe for detecting dead cells [18].

After double staining with these two fluorescent probes, the number and change of living and dead cells can be obtained simultaneously by common fluorescent microscope and flow cytometry, which has been widely used in cell biology.

### Application of fluorescence double staining and Interactive Laser Cytometer technology in tissue engineering research

Chu combined Interactive Laser Cytometer with fluorescence double staining to understand the distribution of cells on polymer carrier. He thought that the advantage of this method over histology and electron microscopy was that the polymer structure would not be distorted, because in the process of tissue treatment, the polymer carrier structure often dissolved in solution and fractured [19].

Interactive Laser Cytometer is especially suitable for the study of anchoring dependent cells [20]. Based on the characteristics of Interactive Laser Cytometer, we quantitatively analyzed the green and red fluorescence intensities of different wavelength. We reported a useful method which can rapidly and quantitatively detect the activity and the proportion of living and dead cells in situ. This method solves the problem that cell growth can’t be observed on opaque carriers, and it doesn’t need to deal with samples as other methods do. In this way the tested engineered tissue can be cultured continuously under the condition of ensuring asepsis, so the growth of each engineered tissue can be dynamically observed. The use of Interactive Laser Cytometer has expanded the scope of application of fluorescence double staining, which is a quantitative method in situ.

## Conclusions

The cultured urethral epithelial cells of male New Zealand young rabbits were capable of extensive expansion in vitro, and can be used for urethral reconstruction. The collagen chitosan composite was more similar to the extracellular matrix of mammalian. Its three-dimensional porous structure had a high area volume ratio, which was conducive to cell adhesion, growth and metabolism. In order to form a tissue-engineered urethra, the cultured urethral epithelial cells were seeded on collagen chitosan composite as scaffolds. CFDA-AM staining showed that the number of living cells was the most when the cells had been cultured on collagen chitosan composite as scaffolds for 7 days. This experiment laid a foundation for the in vivo experiment of tissue-engineered urethra.

## Acknowledgments

We thank the staff at the Plastic Surgery Hospital of Peking Union Medical College and the staff at Tianjin Institute of medical science for assistance.

